# Classification of breast tumours into molecular apocrine, luminal and basal groups based on an explicit biological model

**DOI:** 10.1101/270975

**Authors:** Richard Iggo

**Affiliations:** INSERM U1218, Bergonie Cancer Institute, University of Bordeaux, Bordeaux, France

## Abstract

The gene expression profiles of human breast tumours fall into three main groups that have been called luminal, basal and either HER2-enriched or molecular apocrine. To escape from the circularity of descriptive classifications based purely on gene signatures I describe a biological classification based on a model of the mammary lineage. In this model I propose that the third group is a tumour derived from a mammary hormone-sensing cell that has undergone apocrine metaplasia. I first split tumours into hormone sensing and milk secreting cells based on the expression of transcription factors linked to cell identity (the luminal progenitor split), then split the hormone sensing group into luminal and apocrine groups based on oestrogen receptor activity (the luminal-apocrine split). I show that the luminal-apocrine-basal (LAB) approach can be applied to microarray data (186 tumours) from an EORTC trial and to RNA-seq data from TCGA (674 tumours), and compare results obtained with the LAB and PAM50 approaches. Unlike pure signature-based approaches, classification based on an explicit biological model has the advantage that it is both refutable and capable of meaningful improvement as biological understanding of mammary tumorigenesis improves.

## INTRODUCTION

Early breast cancers are traditionally classified by histology, tumour size, axillary nodal status, grade, Ki67, expression of steroid hormone receptors (oestrogen receptor alpha [ER] and progesterone receptor [PR]) and amplification of human epidermal growth factor receptor 2 (HER2/ERBB2). Together they allow oncologists to select patients for treatment with systemic medical therapies: chemotherapy, drugs targeting ER or oestrogen synthesis, and drugs targeting HER2. Early gene expression microarray studies quickly identified ER positive (luminal), HER2-enriched and ER/HER2 negative (basal or basal-like; for simplicity I will use the term basal) groups (Perou *et al*, 2000; Sorlie *et al*, 2001; Sorlie *et al*, 2003). To make the procedure more robust, the authors subsequently chose a fixed set of 50 genes and defined reference centroids that now form the basis of the widely used “PAM50” classification of breast cancer (Parker *et al*, 2009). In addition to basal and HER2-enriched groups, the PAM50 classification splits the luminal group into low and high proliferation groups (luminal A and B, respectively), and tumours that can not be distinguished from normal tissue are assigned to a “normal” group. An additional “claudin-low” group resembling cell lines that have undergone an epithelial-to-mesenchymal transition (EMT) was added later (Herschkowitz *et al*, 2007; Prat *et al*, 2010).

In 2005 we performed a gene expression study on Affymetrix microarrays that found three main groups: basal, luminal and molecular apocrine (Farmer *et al*, 2005). Our luminal group merges the luminal groups from the PAM50; the basal group is essentially identical in the two classifications. The major disagreement concerns the HER2-enriched and molecular apocrine groups. In the PAM50 it was named “HER2-enriched” because many of the tumours were HER2 amplified. However, about one-third of HER2-enriched tumours are not HER2 amplified, and HER2 amplification is frequently observed in luminal B, occasionally in luminal A and rarely in basal tumours, as recently emphasised in a comprehensive study of HER2 amplification by Daemen and Manning (Daemen & Manning, 2018). In 2005 we suggested that a better name for the HER2-enriched group would be “molecular apocrine” (MA). We based this suggestion on the expression of the androgen receptor (AR) and some genes commonly expressed in prostate cancer, and on the presence of apocrine histological features. We proposed that the tumours in this group were luminal tumours that had undergone apocrine metaplasia. Apocrine metaplasia is a common condition in normal breast tissue in which cells revert from an oestrogen-driven, mammary fate to their ancestral androgen-driven, apocrine fate. Other groups quickly confirmed the existence of a putative androgen-driven group in breast cancer gene expression data (Doane *et al*, 2006; Guedj *et al*, 2012; Lehmann *et al*, 2011). This hypothesis led oncologists to perform clinical trials with anti-androgens (Bonnefoi *et al*, 2016; Gucalp *et al*, 2013; Traina *et al*, 2018). The clinical benefit rate observed in those trials was only 19-25%. While this may seem low, it is comparable to the clinical benefit rate to anti-oestrogen treatment (24-37%) used as second line therapy for ER-positive tumours (Smith & Dowsett, 2003). A further reason for the modest efficacy of anti-androgens in the molecular apocrine trials may have been the inclusion of classic luminal tumours in the groups that received anti-androgens. The key problem leading to misclassification and thus suboptimal treatment is the lack of a precise definition of molecular apocrine tumours.

To go beyond descriptive arguments based on signatures the goal here is to implement a definition for molecular apocrine tumours based on the following explicit biological hypothesis: a molecular apocrine tumour is a hormone sensing cell tumour that has undergone apocrine metaplasia. In the normal mammary gland, luminal progenitors differentiate to form milk secreting cells (M) and hormone sensing cells (H, Fig 1). Recent lineage tracing studies from the Blanpain and Guo groups have identified potential ER-positive stem cells that can maintain the ER-positive lineage through multiple rounds of grafting (Van Keymeulen *et al*, 2017; Wang *et al*, 2017). This ER-positive stem cell (marked “H” in Fig 1) is potentially the cell of origin of human luminal and molecular tumours. Upon malignant transformation, I propose that ER-positive stem cells occasionally undergo apocrine metaplasia, lose ER expression and give rise to molecular apocrine tumours. Lineage tracing has also identified an ER-negative Notch1-derived cell that is potentially the cell of origin of human basal tumours (marked “M” in Fig 1) (Rodilla *et al*, 2015). Unlike the classic Lim model (Lim *et al*, 2009), the model in Fig 1 places basal tumours distal to luminal progenitors on the secretory branch because luminal progenitors express ER and ELF5, whereas basal tumours are rigorously ER-negative. The model explicitly states that molecular apocrine tumours are a subtype of hormone sensing cell tumour (Fig 1).

**Figure 1.**
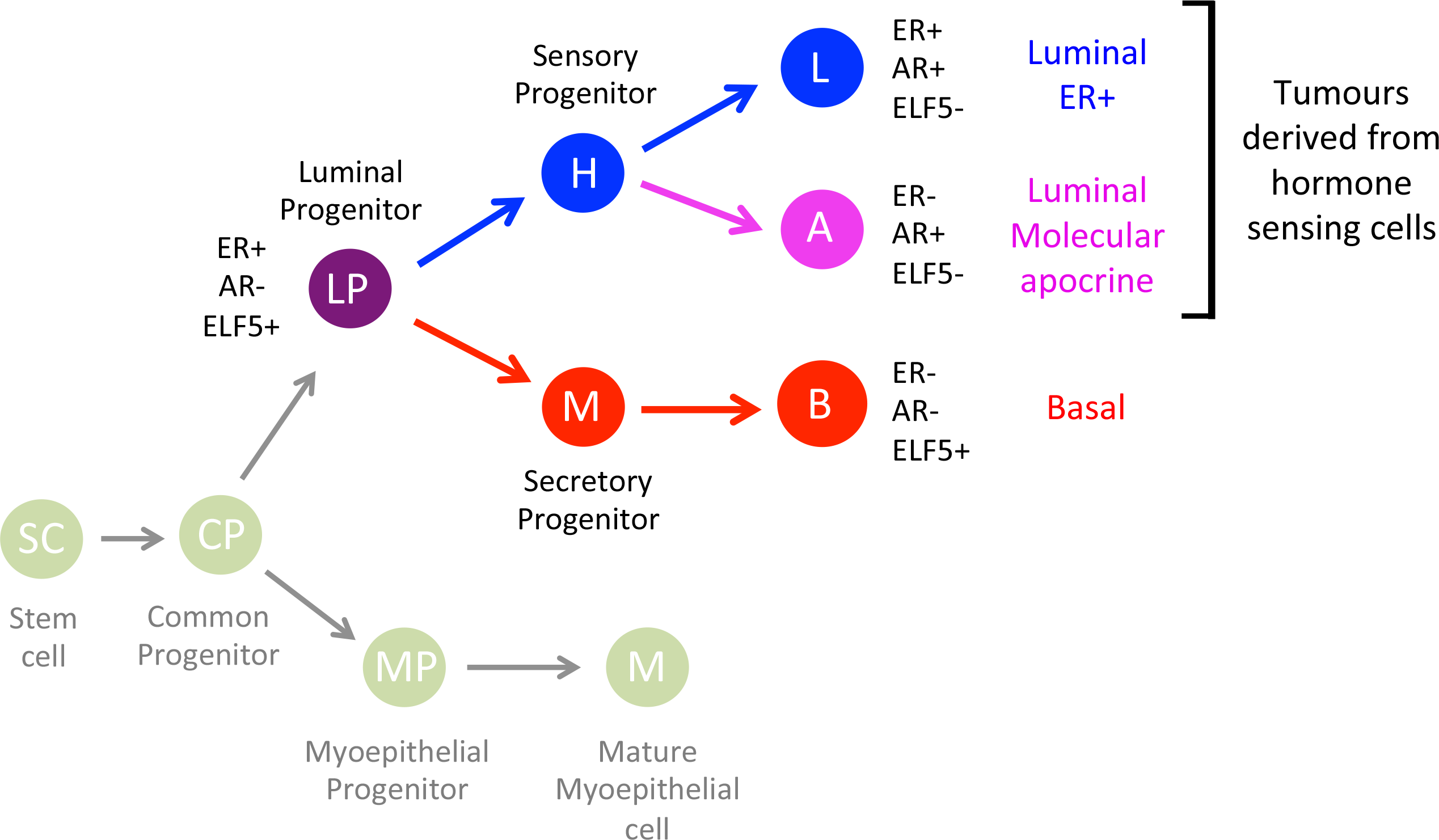
Mammary lineage diagram. Luminal progenitors give rise to hormone-sensing and milk-secreting lineages. ELF5 and ER are transcription factors that act as master regulators for these lineages. Lineage tracing shows that the sensory and secretory progenitors marked H (hormone) and M (milk) can be serially transplanted indicating that they have the properties of stem cells for their respective lineages. These lineage-specific stem cells are potentially the cells of origin of most human breast tumours. The nomenclature is confusing because all cells derived from luminal progenitors (luminal, molecular apocrine and basal) are anatomically luminal (ie, line the inner layer of the ducts) but the term luminal is sometimes used to mean only ER+ luminal tumour cells (as in “luminal A/B”) and sometimes to mean any tumour cells specialised for hormone sensing (ER+ and molecular apocrine). Note that AR is expressed by all cells in the hormone-sensing lineage. The term luminal AR+ (LAR) to mean molecular apocrine is thus particularly confusing because luminal ER+ tumours are luminal and AR+ but they are not included in the LAR group. Most mouse mammary tumours are derived from the green cells in the diagram (anatomically basal cells) and form tumour types rarely seen by human pathologists.

The first step in the classification (the “luminal progenitor split”) is based on expression of transcription factors that are known or suspected to play a role in defining secretory or sensory cell identity. The master regulator for the secretory lineage is ELF5 (Kalyuga *et al*, 2012; Oakes *et al*, 2008; Zhou *et al*, 2005), and for the sensory lineage it is ER (Curtis Hewitt *et al*, 2000; Mallepell *et al*, 2006), but they cooperate with other transcription factors that are themselves closely linked to cell fate. The first step in the classification is thus to split tumours into sensory cell tumours expressing ESR1, AR, FOXA1, TOX3, SPDEF, GATA3, MYB, MSX2, TFAP2B and ESRRG; and secretory cell tumours expressing FOXC1, BCL11A, ELF5, KLF5, VGLL1, NFIB, ID4, SOX10 and EN1. These transcription factors show strongly bimodal expression in breast tumours allowing clean separation of the two cell types. They were selected based on published studies on mammary development, hormonal signalling and differential expression in normal and transformed mammary epithelial cells. MSX2 and BCL11A are directly implicated in defining mammary cell identity at the earliest stages of mammary gland development (Howard & Ashworth, 2006; Khaled *et al*, 2015). ELF5, the master regulator of lactation, is necessary for the formation of milk secreting cells (Oakes *et al*, 2008; Zhou *et al*, 2005). ER, AR, FOXA1 and GATA3 play crucial roles in the transcriptional response to androgens and oestrogens (Carroll *et al*, 2005; Robinson *et al*, 2011). Some factors, like SPDEF, TOX3 and KLF5, are probably responsible for specific subprograms that contribute to mammary cell function (Oishi *et al*, 2008; Raap *et al*, 2018; Seksenyan *et al*, 2015). Others, like VGLL1, EN1 and SOX10, are components of classic developmental pathways (Loomis *et al*, 1996; Tsurusaki *et al*, 2014; Vaudin *et al*, 1999). Translocation of MYB to NFIB perturbs mammary lineage decisions, resulting in the formation of adenoid cystic tumours with distinct luminal and myoepithelial tumour cell populations (Persson *et al*, 2009). Many of these genes are known to be differentially expressed in purified mammary cell subsets (Asselin-Labat *et al*, 2007; Kendrick *et al*, 2008).

The second step is to split sensory cell tumours into classic ER+ luminal tumours and molecular apocrine tumours. The critical underlying event is apocrine metaplasia accompanied by loss of ER expression but the biological mechanism is not currently understood; the working model is that an epigenetic event switches the cell between mammary and apocrine programs. The difference between these programs is that the former is regulated by oestrogens and the latter by androgens. Since AR itself is expressed by both luminal and apocrine cells it can not be used to make the distinction. Pending greater understanding of the underlying epigenetic event the most useful marker is ER. I therefore created luminal and apocrine scores based on genes with the strongest positive and negative correlation with ESR1, respectively, using gene expression data (Farmer 2005; Farmer 2009) generated from a subset of tumours from the EORTC 10994 trial (Bonnefoi *et al*, 2011). Only sensory cell tumours (luminal and molecular apocrine) were included in the subset used for this comparison to avoid selecting genes that simply distinguish between basal and luminal tumours. I suspect that genes showing the strongest negative correlation are induced by AR in cells that have undergone apocrine metaplasia but the lack of good cell or animal models makes this difficult to prove. Unlike models based purely on signatures, this model is both falsifiable and likely to improve as biological understanding of apocrine metaplasia increases, in particular when cell culture models are developed.

This paper contains 1. scripts to classify breast tumours, 2. application of the approach to an independent dataset, 3. comparison with the PAM50 classification, and 4. comparison with two-gene predictors. Since the approach leads to three classes (Luminal, molecular Apocrine and Basal) I refer to it as the LAB classification.

## METHODS

The EORTC 10994 phase III clinical trial tested whether p53 mutant tumours respond better to anthracycline-based chemotherapy that includes taxanes (Bonnefoi *et al*, 2011). The trial was registered with ClinicalTrials.gov number NCT00017095 and approved by national and/or local ethics committees in all participating centres. Before registration, all patients signed an informed consent for the trial and for research on tumour samples. Microarray data from 186 tumours included in our previous gene expression studies (Farmer *et al*, 2009; Farmer *et al*, 2005), was downloaded from the NCBI GEO database entries with accession numbers GSE1561 and GSE6861. The batches and sample names in the GSE1561 study (49 samples) and GSE6861 study (161 samples) are given in Sup Table 1.

The EORTC 10994 trial enrolled patients with T2-T4 M0 tumours with ≥20% tumour cells in the pretreatment biopsy. The RNA was extracted from a 200 um thickness of a 14G needle biopsy. The small size of the samples reduced the scope for tumour content to drift between the section examined by the pathologist and the material tested on the microarray. This may explain the rather low normal tissue contamination compared to studies using surgical samples. The tumours arrayed are not a random selection of tumours in the EORTC 10994 clinical study; instead they contain more ER-tumours leading to a more equal representation of luminal, molecular apocrine and basal tumours than in studies like TCGA (TCGA, 2012) which are overwhelmingly ER+.

To reduce overfitting to a single chip type, the data from the two EORTC 10994 microarray studies (GSE1561 and GSE6861) were pooled. GSE1561 was performed on Affymetrix U133A chips; GSE6861 was performed on Affymetrix X3P chips. The probesets in the former lie within 600 bp of the polyadenylation site, those in the latter within 300 bp of the polyadenylation site. The two datasets were combined with COMBAT (R sva package, Johnson *et al*, 2007) after quantile normalisation of the chip types separately with rma (R affy package, Gautier *et al*, 2004). Since all such procedures expect the same spectrum of variation in each batch, COMBAT was only used to normalise for the difference in chip type. Before combining the datasets very short probesets were removed from the X3P data since they cluster together regardless of the ostensible target gene. Clustering shows that samples hybridised to different chips cluster together after normalisation with COMBAT (Sup Fig 1 and Sup Data 6).

To test whether the classification procedure could be applied to RNA-seq data a prenormalised table of gene expression values from the TCGA molecular portraits study (TCGA, 2012) was downloaded from http://research-pub.gene.com/HER2pancancer (Daemen & Manning, 2018) and 674 stage II-III tumours were analysed. To determine whether the LAB classification produces broadly similar results to the PAM50 classification (Perou *et al*, 2010) the EORTC and TCGA tumours were classified with a script provided by Dr Joel Parker (https://genome.unc.edu/pubsup/breastGEO/PAM50.zip) using the default parameters. Median files for the EORTC and TCGA matrices were used to reduce platform bias as described (Perou *et al*, 2010).

Hierarchical clustering was performed in Cluster with median centring, uncentred correlation distance and centroid linkage in Cluster (de Hoon *et al*, 2004; Eisen *et al*, 1998). Heatmaps were created in Treeview (Eisen *et al*, 1998; Saldanha, 2004). The bars showing the LAB and PAM50 class were created in the heatmap.plus R package (v1.3, Allen Day 2012, cran repository). All other procedures used to process the data were performed as described in the supplementary data (Sup Data files 1-5).

## RESULTS

### The LAB classification of breast tumours

The first step in the LAB classification of breast tumours is to split tumours into sensory cell (H) versus secretory cell (M) tumours (Fig1) based on the expression of transcription factors known or suspected to play an important role in defining mammary cell identity. I defined sensory and secretory scores as the mean expression values of the respective cell identity transcription factors after scaling. The EORTC tumours form two clearly distinct clusters based on the sensory and secretory scores (Fig 2a), yielding a strongly bimodal luminal progenitor score after 45° rotation of the data (Fig 2b&c). I modelled the luminal progenitor scores as a mixture of normal distributions and defined a cut-off to distinguish the two tumour types (Fig 2d) and normalised the scores to place the peaks at -1 and 1 (Fig 2e).

**Figure 2.**
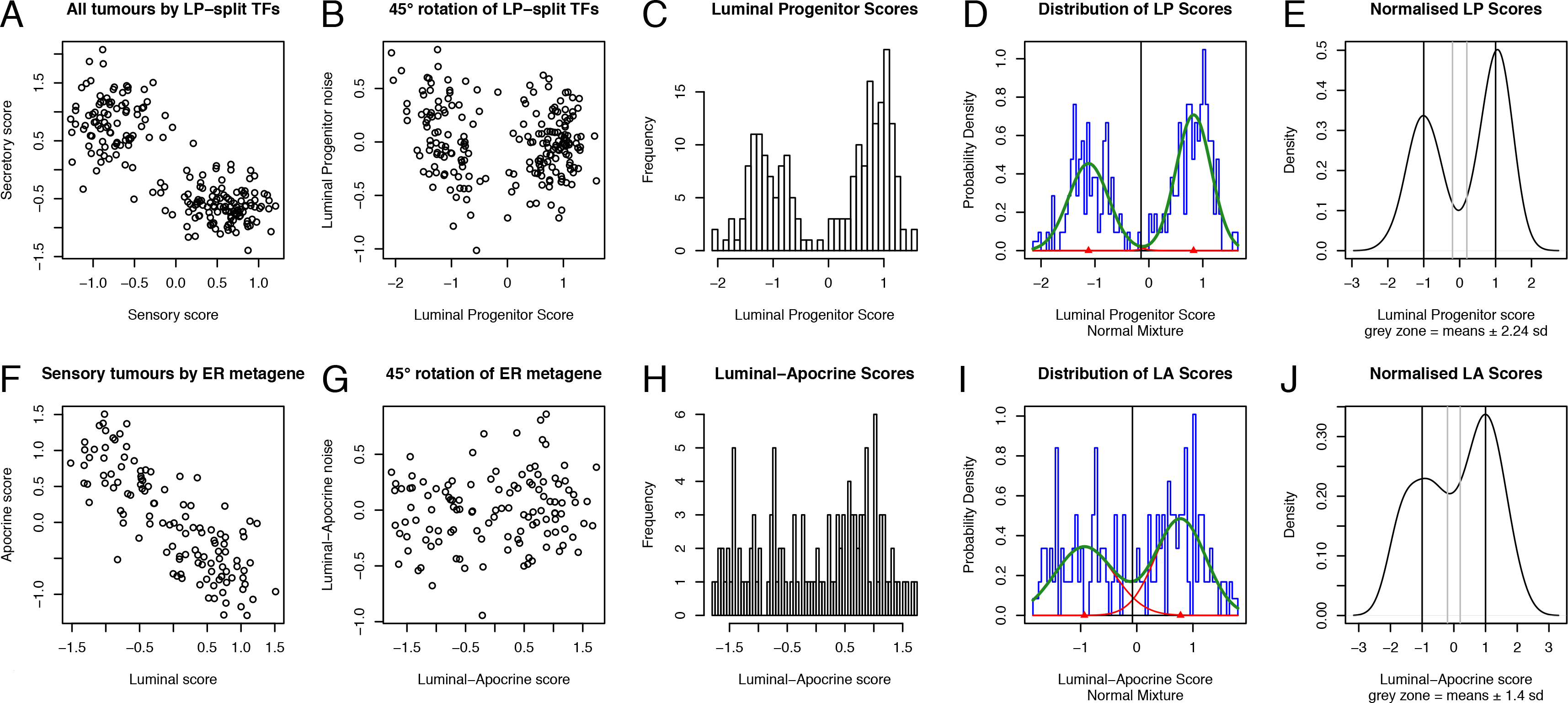
Processing steps for the EORTC data. The EORTC tumours (210 samples) were split into sensory and secretory lineages in the upper panels (A-E) then the sensory tumours were split into luminal and molecular apocrine tumours in the lower panels (F-G). A, The sensory and secretory scores are the mean scaled expression values for transcription factors that split luminal progenitors into sensory and secretory cells (LP-split TFs). B, 45° rotation to place the major variation on one axis. C, Histogram of the luminal progenitor (LP) scores (x-axis from panel B). D, Fitted mixture of normal distributions of LP scores. E, LP scores normalised by fixing the means of the normal distributions at −1 and 1, with a cut-off at 0. The grey vertical lines at −0.2 and 0.2 define the grey zone in which tumour class is called as unknown, with the distance to the respective means indicated as a multiple of the average standard deviation of the normal distributions. The upper peak in E contains the sensory cell tumours; these tumours were split into luminal and molecular apocrine tumours in panels F-J. F, The luminal and apocrine scores are the mean scaled expression values of 15 genes positively correlated with ER (luminal score) and 15 genes negatively correlated with ER (apocrine score). G-J, The procedures in the upper panels were applied to the Luminal and Apocrine scores to classify the sensory tumours into luminal and molecular apocrine tumours.

Since tumours lying exactly at the cut-off can not sensibly be assigned to any group, tumours scoring between -0.2 and 0.2 were called as “unknown”. The lower peak defines basal tumours in the LAB classification (the red branch in Fig 1). To give some insight into what the -0.2 and 0.2 boundaries might mean they are expressed in Fig 2e as a multiple of the standard deviation in the mixture model. For the luminal progenitor scores the value in the EORTC data was 2.24, indicating that about 99% of tumours classified as basal are likely to lie closer to the secretory cell mean than to the sensory cell mean. The value depends on the quality of the estimate of the standard deviation in the mixture model so it should be taken as a rough guide rather than a precise measure.

To separate luminal from molecular apocrine tumours, 30 genes were selected based on correlation with ESR1 expression, half showing positive (luminal) and half negative (molecular apocrine) correlation. The genes are listed in Table 1. The classification procedure was then repeated exactly as for the luminal progenitor scores (Fig 2, lower panels). The luminal-apocrine score is less bimodal than the luminal progenitor score, leading to greater overlap of the luminal and apocrine distributions. The distance to the grey zone is consequently only 1.4 times the standard deviation, indicating that only about 90% of tumours are likely to be correctly classified. The LAB class assignments are shown in Table 2: 29% luminal, 23% molecular apocrine, 42% basal and 6% unknown. The bias towards molecular apocrine and basal tumours is expected because the samples chosen for the EORTC microarray studies were deliberately enriched in ER-negative tumours.

**Table 1.** LA genes. These genes were used to split luminal tumours from molecular apocrine tumours in the lower panels in Figs 2 and 3. The Human Genome Organisation Gene Nomenclature Committee gene symbol is shown. The genes are shown in descending order of positive (Luminal) or negative (Molecular apocrine) correlation.

| Luminal | Molecular apocrine |
| --- | --- |
| ESR1 | CLCA2 |
| CA12 | KMO |
| BCL2 | PAPSS2 |
| GFRA1 | PSAT1 |
| GREB1 | KYNU |
| FAM134B | KRT7 |
| IGF1R | RARRES1 |
| NPY1R | AKR1B10 |
| ANXA9 | SOX11 |
| SERPINA5 | TFAP2B |
| SCCPDH | PERP |
| IRS1 | HSD17B2 |
| ABAT | DKK1 |
| SERPINA3 | LIMCH1 |
| MTL5 | IL8 |

**Table 2.** LAB vs PAM50 classification of EORTC tumours. Contingency table comparing the results of classification with the LAB and PAM50 algorithms. MA, molecular apocrine; LumA, luminal A; LumB, luminal B.

|  |  | LAB classification |  |  |  |
| --- | --- | --- | --- | --- | --- |
|  |  | Luminal | MA | Basal | unknown |
| PAM50<br>classification | LumA | 36 | 9 | 0 | 4 |
|  | LumB | 23 | 0 | 0 | 2 |
|  | Her2 | 1 | 34 | 0 | 6 |
|  | Basal | 0 | 3 | 83 | 0 |
|  | Normal | 1 | 2 | 5 | 1 |

### Application to an independent dataset

To test whether the LAB classification can be applied to other types of gene expression data the procedure was repeated on RNA-seq data from The Cancer Genome Atlas (TCGA, 2012). Fig 3 shows the result of analysing 674 stage II-III tumours. The figure is strikingly similar to Fig 2, apart from the much higher fraction of luminal tumours in the TCGA dataset (Table 3: 70% luminal, 7% molecular apocrine, 17% basal, 5% unknown). The high percentage of ER-positive tumours in the TCGA dataset is typical of breast cancer in the general population. It is difficult to give a precise figure for the expected fraction of molecular apocrine tumours in an unselected population but a figure of 7% is plausible. For example, a large French microarray study recently classified 11% of tumours as molecular apocrine (Guedj *et al*, 2012). Hierarchical clustering shows the expected patterns: a homogeneous basal group, a small core molecular apocrine group and a large luminal group (Sup Fig 2; the red, pink and blue bars at the top of the heatmap show the LAB and PAM50 classifications). I conclude that it is technically feasible to transfer the LAB classification across platforms and studies.

**Figure 3.**
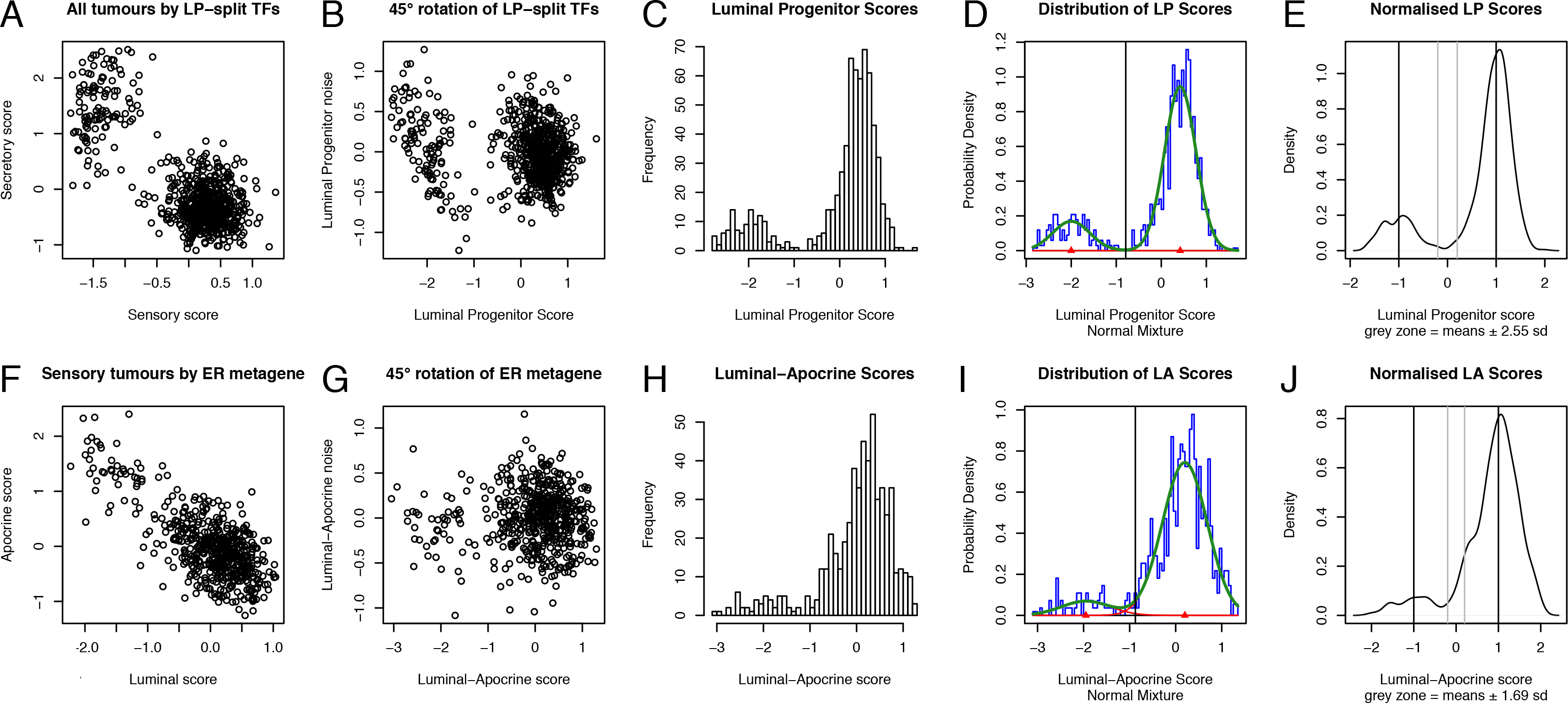
Processing steps for the TCGA data. The TCGA tumours (674 samples) were classified with the same two-step classification scheme used for the EORTC tumours. See Fig 2 for details.

**Table 3.** LAB vs PAM50 classification of TCGA tumours. Contingency table comparing the results of classification with the LAB and PAM50 algorithms. MA, molecular apocrine; LumA, luminal A; LumB, luminal B.

|  |  | LAB classification |  |  |  |
| --- | --- | --- | --- | --- | --- |
|  |  | Luminal | MA | Basal | unknown |
| PAM50<br>classification | LumA | 330 | 5 | 0 | 16 |
|  | LumB | 140 | 4 | 0 | 11 |
|  | Her2 | 4 | 39 | 0 | 6 |
|  | Basal | 0 | 1 | 112 | 2 |
|  | Normal | 0 | 0 | 4 | 0 |

### Comparison with the PAM50 classification

For the EORTC tumours, 43 of the PAM50 genes were successfully mapped to the Affymetrix dataset (the missing genes are not present on the U133A chip). Nineteen tumours were incomparable (classified as Normal by PAM50, or unknown by LAB), leaving 191 samples to compare (Table 2). There was excellent agreement for the luminal and basal tumours: 98% (59/60) of tumours classified as luminal by LAB were classified as luminal A or B by PAM50, and 100% (83/83) of tumours classified as basal by LAB were classified as basal by PAM50. In contrast, only 74% (34/46) of those classified as molecular apocrine by LAB were classified as HER2-enriched by PAM50. Most of the disagreements were molecular apocrine called as luminal A by PAM50. Some of the difference can probably be explained by the seven PAM50 genes lost to cross-platform mapping.

For the TCGA tumours, all of the PAM50 genes were successfully mapped to the RNA-seq dataset. Thirty-nine tumours were incomparable (classified as Normal by PAM50, or unknown by LAB), leaving 635 tumours to compare. There was excellent agreement for the luminal and basal tumours: 99% (470/474) of tumours classified as luminal by LAB were classified as luminal A or B by PAM50, and 100% (112/112) of tumours classified as basal by LAB were classified as basal by PAM50 Table 3). In contrast, only 80% (39/49) of tumours classified as molecular apocrine by LAB were classified as HER2-enriched by PAM50. All but one of the disagreements between the two classifications were HER2-enriched tumours called luminal by LAB, or molecular apocrine called as luminal A or B by PAM50. Heatmaps after hierarchical clustering show visually the strong overall agreement between the LAB and PAM50 assignments (Sup Figs 1&2, the assignments are shown in red, pink and blue in the colour bars at the top of the heatmaps). Unfortunately, the disagreements are concentrated in the luminal/apocrine populations, where misclassification has important implications for hormonal therapy.

### Comparison with two-gene predictors

Pathologists commonly use ER, AR and HER2/ERBB2 to identify molecular apocrine tumours. To explore the potential utility of these genes to define molecular apocrine status the expression of ER vs ERBB2 and ER vs AR in the EORTC and TCGA datasets was plotted (Fig 4, left and centre panels). Molecular apocrine tumours are expected to reside in the upper left quadrants with low ER expression and high ERBB2 or AR expression. The individual points are labelled with the LAB classification in Fig 4a&c and with the PAM50 classification in Fig 4b&d. The ER vs ERBB2 plots confirm that ERBB2 alone is not a good way to predict molecular apocrine status, with many ERBB2-overexpressing tumours in the upper right (luminal) quadrant. Another drawback of ERBB2 is that some molecular apocrine tumours are buried in the lower left (basal) quadrant. The ER vs AR plots likewise show that AR alone is not a good way to classify the tumours, with many luminal tumours having AR expression overlapping with that of molecular apocrine and basal tumours. FOXA1 is more bimodal than AR (Fig 5). It produces three distinct clusters that correspond well to the LAB classification (Fig 4a&c), with molecular apocrine tumours in the upper left quadrant, luminal tumours in the upper right quadrant and basal tumours in the lower left quadrant. The lower right quadrant is empty, as expected because ER function requires FOXA1. On the basis of this analysis, ER and FOXA1 could potentially form the basis for an IHC test for molecular apocrine tumours.

**Figure 4.**
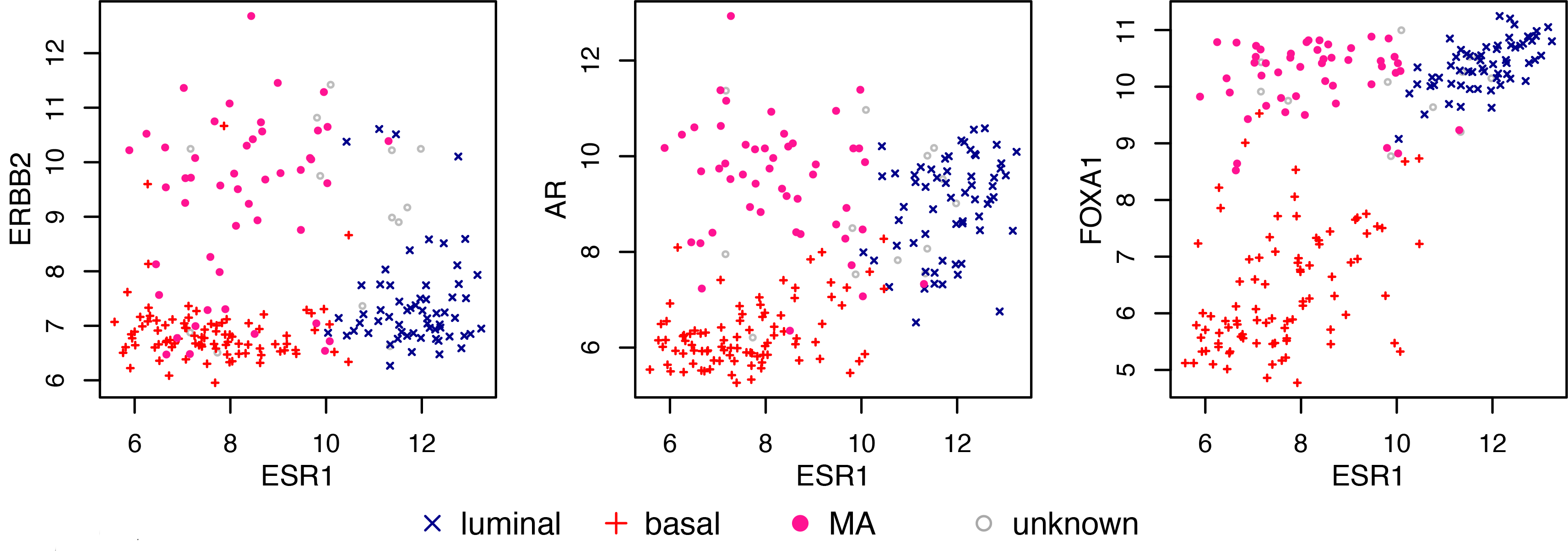
EORTC and TCGA tumours showing ESR1 expression plotted against ERBB2, AR and FOXA1 expression for each tumour. The points are coloured according to the LAB classification in a&c and according to the PAM50 classification in b&d. a. LAB classification of EORTC tumours, b. PAM50 classification of EORTC tumours, c. LAB classification of TCGA tumours, d. PAM50 classification of TCGA tumours. The units on the x and y axes are arbitrary gene expression values: log2 rma-normalised intensities for the EORTC Affymetrix data; log2 reads per kilobase of exon model per million mapped reads normalised by size factor [log2 nRPKM+1] for the TCGA data (Daemen & Manning, 2018).

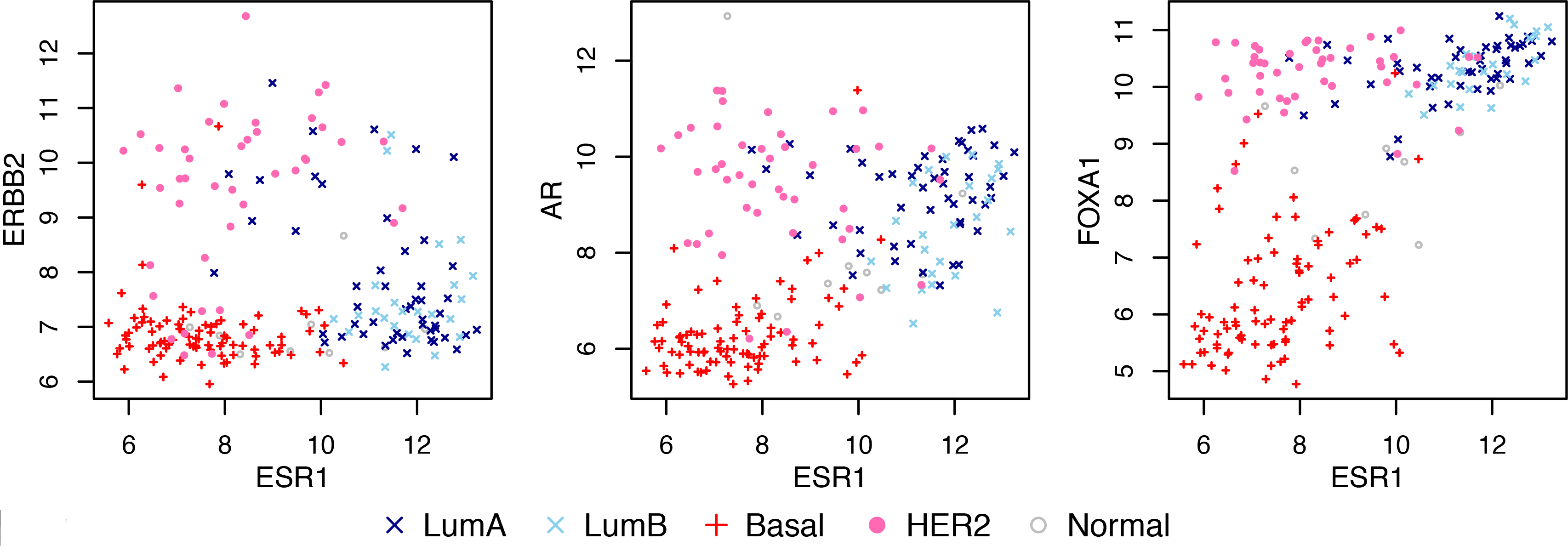

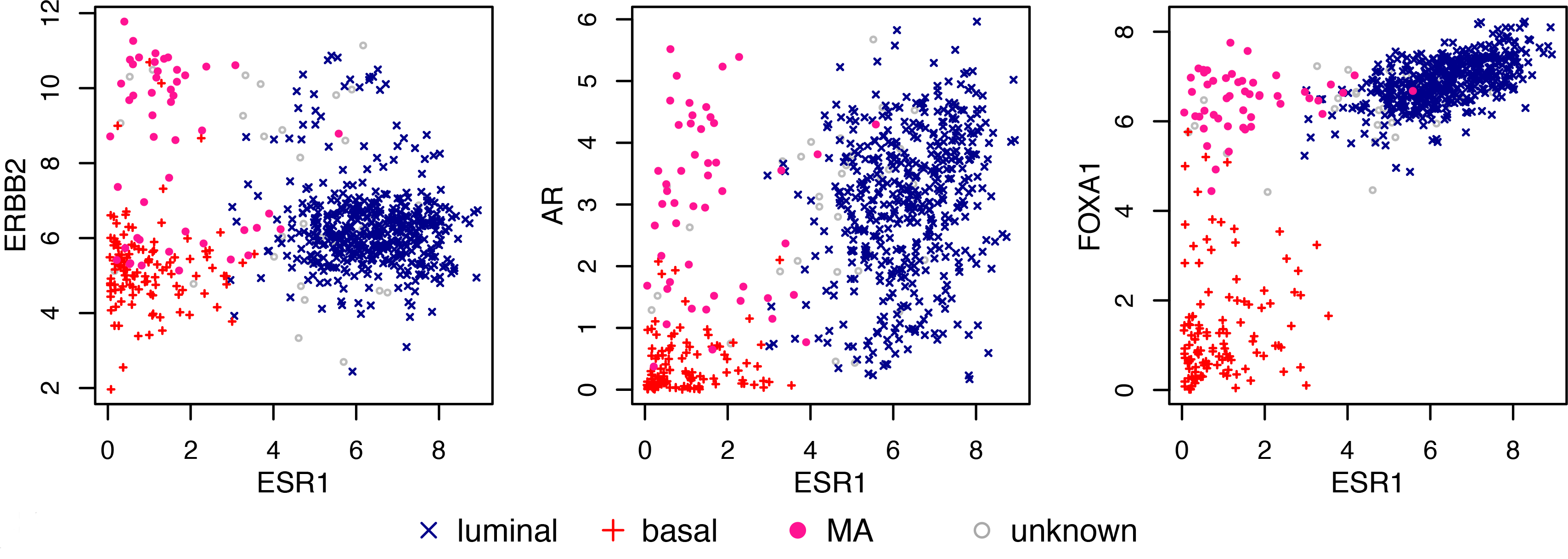

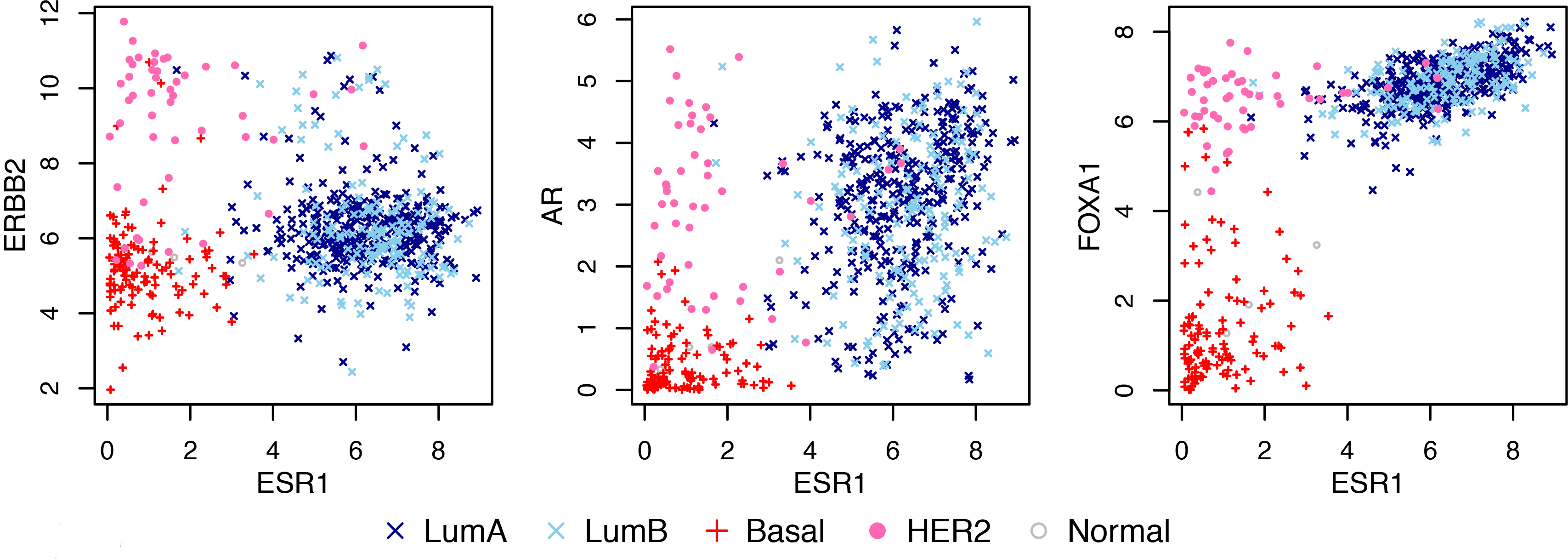

**Figure 5.**
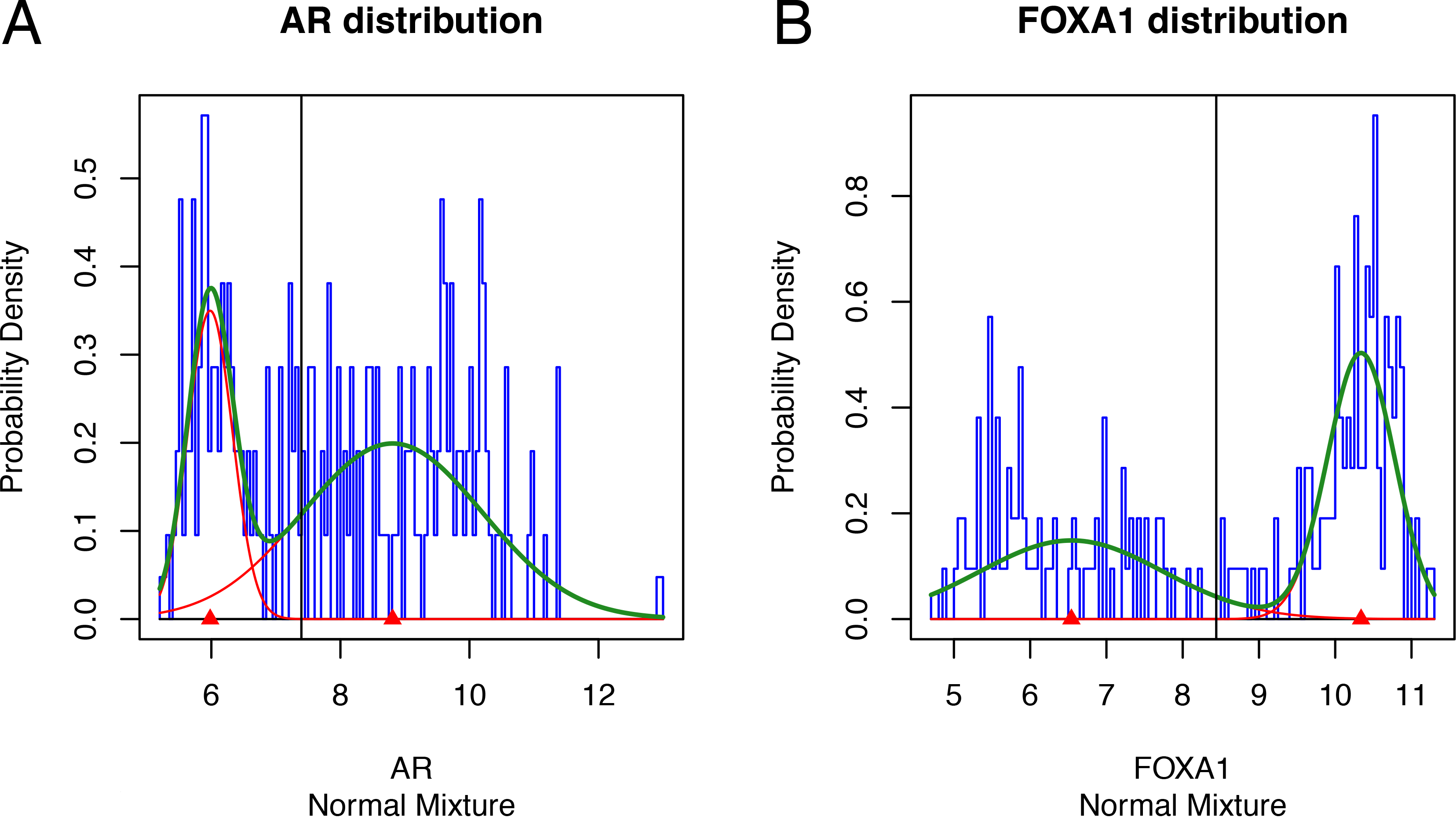
Bimodality of AR and FOXA1 expression in the EORTC tumours. A, AR has a broad distribution that is difficult to resolve into two separate peaks. b, FOXA1 has a bimodal expression pattern. The FOXA1-high peak contains the molecular apocrine and luminal tumours; the FOXA1-low peak contains the basal tumours. AR can not be used to separate the tumours in the same way because there is no natural cut-off between the groups.

## DISCUSSION

I have shown that the three main types of breast cancer can be identified with gene expression data on the basis of the lineage model shown in Figure 1. One advantage of rooting a classification in biology is that it is falsifiable. Another is that it can improve as understanding of mammary biology improves. This is particularly true of the luminal-apocrine split which currently rests on loss of ER activity rather than a deep mechanistic understanding of apocrine metaplasia. This reflects the genuine difficulty we currently face in trying to understand what it means to be a molecular apocrine tumour. Metaplasia is a change in cell identity that may not be caused by any driver mutation. Instead, it probably results from epigenetic changes that lock in self-sustaining patterns of transcription factor expression through positive feedback. It is not obvious what these factors are, but recent progress in breast cancer modelling is likely to provide true molecular apocrine models that may shed light on the mechanism of apocrine metaplasia (Sachs *et al*, 2018; Verbeke *et al*, 2014). Positive feedback loops are broken by external signals when cells change identity. The critical signal regulating the luminal-apocrine transition in mammary epithelial cells is probably transmitted by the ERBB2/ERBB3 pathway. Predisposition of sensor cells to apocrine metaplasia by ERBB2 amplification would then explain the very strong overlap of ERBB2 amplification with molecular apocrine differentiation.

Transferring signatures across platforms is notoriously unreliable, partly because of batch effects and overfitting to the starting platform (Kratz & Carninci, 2014) but also because differences in the spectrum of tumours present in individual studies can introduce unwanted bias during normalisation (Zhao *et al*, 2015). Approaches based on biology should be less sensitive to these issues. Indeed, it was surprisingly easy to transfer the LAB classification to RNA-seq data. It should be noted, however, that the LAB algorithm automatically reduces differences between datasets in a manner that would not meet the criteria for a universal clinical test. In particular, bimodal decomposition of the distribution of the scores to find the cut-off is sensitive to the composition of the dataset. Once the thresholds for a particular platform have been defined it should be easy to apply the algorithm on a case-by-case basis but that was not the goal of this study.

To verify that the LAB approach generates results that are broadly consistent with existing approaches it was compared to the widely used PAM50 test (Parker *et al*, 2009). In considering the results it is important to note that the two approaches have different objectives. The PAM50 classifier encompasses proliferation, ER status, HER2 status and normal tissue contamination. The focus of the LAB classifier is to separate tumours based on an underlying biological model, not to measure proliferation or normal tissue contamination. There was excellent agreement in both the EORTC and TCGA datasets over the identification of luminal and basal tumours. Where the two classifications were expected to disagree is over the HER2-enriched and molecular apocrine groups, and this is indeed what happened. The difficulty of distinguishing luminal B from HER2-enriched tumours in the PAM50 classification has been widely commented on (Farmer *et al*, 2005; Mackay *et al*, 2011; Weigelt *et al*, 2011), including by the original authors of the PAM50 classification (Parker *et al*, 2009). In some cases the tumours in the luminal cluster called as HER2-enriched had increased ERBB2 expression. This is broadly expected because the main determinant of HER2-enriched status in the PAM50 classification is expression of two genes in the ERBB2 amplicon (ERBB2 and GRB7). Since ERBB2 is commonly amplified in luminal ER-positive tumours, it is inevitable that some luminal tumours will be misclassified as HER2-enriched. The root cause of the problem is that gene expression arrays measure phenotype not genotype. ERBB2 amplification is a DNA change (ie, genotype) that is seen in many tumour types that do not have a molecular apocrine phenotype (Daemen & Manning, 2018). There is an excellent correlation between ERBB2 amplification and ERBB2 expression, but it contributes only a small part to the molecular apocrine phenotype. This, and the fact that ERBB2 is amplified in both luminal and molecular apocrine tumours, make it an unreliable marker for molecular apocrine tumours. This explains why ERBB2 is not present in the gene list used for the luminal-apocrine split.

Unlike the PAM50, the LAB classification contains an “unknown” category. This reflects the reality that some tumours resemble intermediate states, perhaps because some oncogenes induce plasticity that allows tumour cells to change identity (Koren *et al*, 2015; Van Keymeulen *et al*, 2015). Whether to assign all tumours to a tumour class, as in the PAM50 classification, or to accept that some can not be classified, as in the LAB classification, is a philosophical question. In the PAM50 a tumour is given the class of the reference centroid to which it shows the highest correlation. An extreme example would be a tumour with a correlation of 0.50 to the normal centroid and a correlation of 0.51 to the luminal A centroid. In that case the coefficient of determination (r^2^) differs by only 1% between the two assignments, meaning the decision is based on only 1% of the information in the profiles. Clinicians faced with binary decisions (treatment A vs treatment B) may prefer always to receive a tumour class, even when they know it is based on imperfect information. If that were the goal it would be easy to remove the unknown group in the LAB classification.

Outside clinical trials, pathologists rarely use microarray or RNA-seq technology to classify tumours. They use IHC for ER, PR and Ki-67 to identify oestrogen-dependent, low proliferation, good prognosis tumours (luminal A); and HER2 IHC or FISH to identify HER2-dependent tumours. Molecular apocrine tumours are a particular problem for them. In principle molecular apocrine tumours are stimulated by androgens whereas luminal tumours are inhibited by androgens (Hickey *et al*, 2012). This means wrong diagnosis could result in patients receiving treatment that stimulated tumour growth. Since the model is that molecular apocrine tumours are driven by AR it is tempting to use AR alone to identify molecular apocrine tumours but this will always fail because classic ER-positive luminal tumours also express AR. The term LAR (luminal AR+, Lehmann *et al*, 2011) for molecular apocrine tumours feeds this misunderstanding. TNBC (triple negative breast cancer) is a clinical term based on IHC for ER, PR and HER2, supplemented with FISH for HER2 in some cases, but it is commonly equated with basal tumours. The lineage diagram in Fig 1 shows why it makes no biological sense to think of LAR tumours as a subset of TNBC or basal tumours: luminal and molecular apocrine tumours are derived from hormone-sensing cells not secretory cells. In principle, TNBC tumours should all be ER-negative but this seems not to be the case, presumably because IHC does not always perfectly capture ER/PR status. Indeed, 82% of the tumours in the LAR group in the Lehmann TNBC study and 63% of the tumours in a subsequent TNBC study from Houston were classified as luminal A or B (Burstein *et al*, 2015; Lehmann *et al*, 2011). The distribution of AR gene expression values in breast tumours is rather broad (Figs 4 and 5), and IHC is less reliable than gene expression assays for quantification of AR level (Lehmann-Che *et al*, 2013), further increasing the difficulty for pathologists. HER2 status is part of the definition of TNBC but it is irrelevant to the definition of molecular apocrine tumours, adding to the confusion. It is important to avoid these misunderstandings when selecting patients for trials with anti-androgens because inclusion of patients with classic oestrogen-dependent luminal tumours would reduce the power to detect a therapeutic response of molecular apocrine tumours to anti-androgens. Based on the analysis in Figs 4&5, IHC for FOXA1 could potentially be used to improve the identification of molecular apocrine tumours in routine diagnosis, but pathologists are unlikely to abandon the use of AR, not least because it is the target of anti-androgens. Ultimately, I hope that rooting the classification of molecular apocrine tumours in mammary gland biology will help to reduce the misunderstandings that surround the definition of molecular apocrine tumours and thereby facilitate the development of treatments for this poor prognosis subtype of breast cancer.

## Acknowledgements

I thank the Fondation pour la lutte contre le cancer et pour des recherches medico-biologiques, the SIRIC BRIO (Grant INCa-DGOS-INSERM 6046) and French Cancer League Comité des Landes for financial support. I thank Anneleen Daemen and Hervé Bonnefoi for critically reading and providing excellent feedback on the manuscript.

---

**Sup Fig 1.**
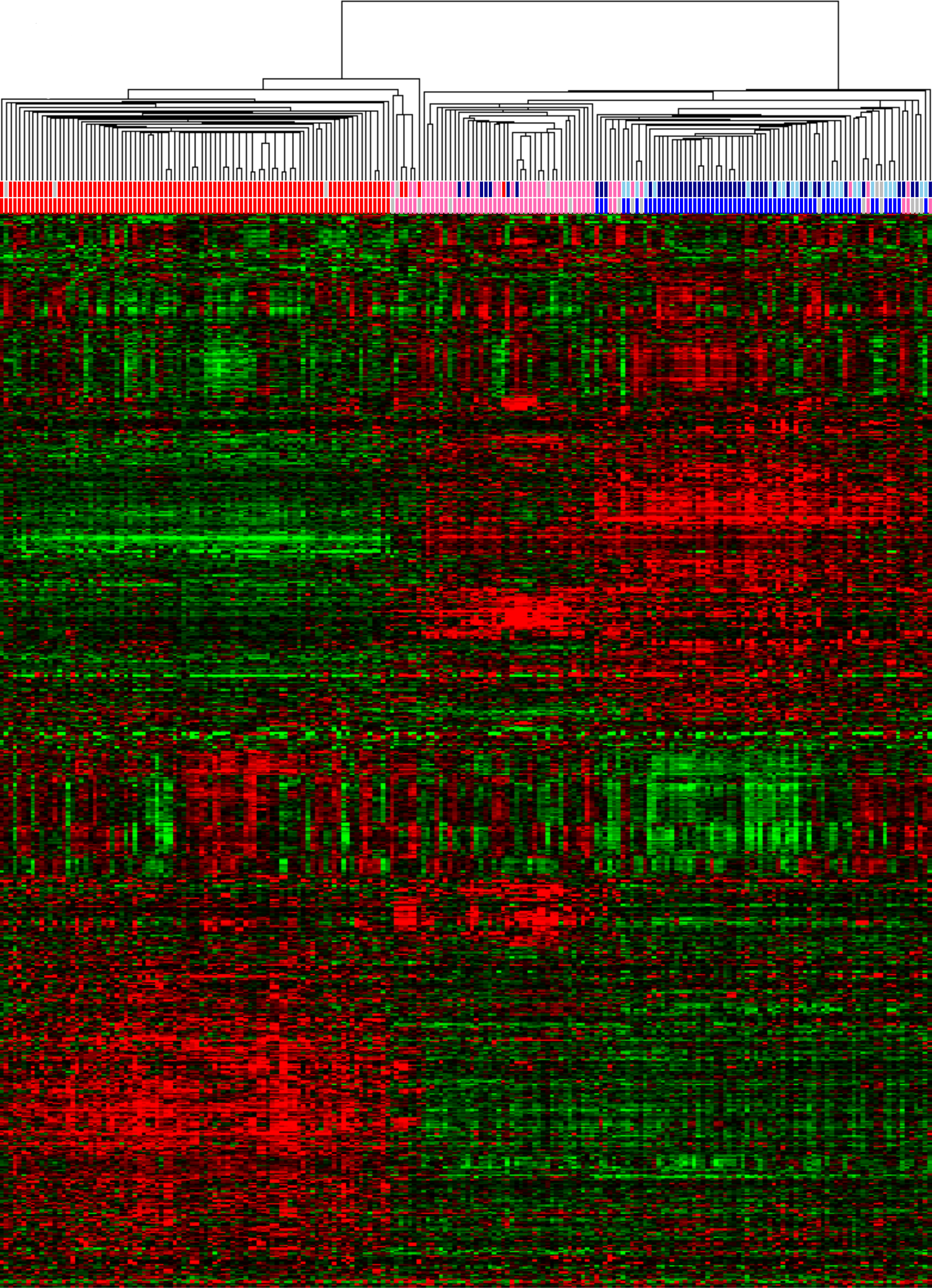
EORTC heatmap. Hierarchical clustering of the 676 most variable genes in the EORTC dataset. The upper row in the colour bar at the top of the heatmap shows the PAM50 classification (basal, red; HER2, pink; luminal A, dark blue; luminal B, light blue; normal, grey). The lower row in the colour bar at the top of the heatmap shows the LAB classification (basal, red; molecular apocrine, pink; luminal, blue; unknown, grey). To view the individual gene and tumour annotations please use the cdt file in Sup Data 6 or use the script in Sup Data 1 to regenerate the full input files for Cluster.

**Sup Fig 2.**
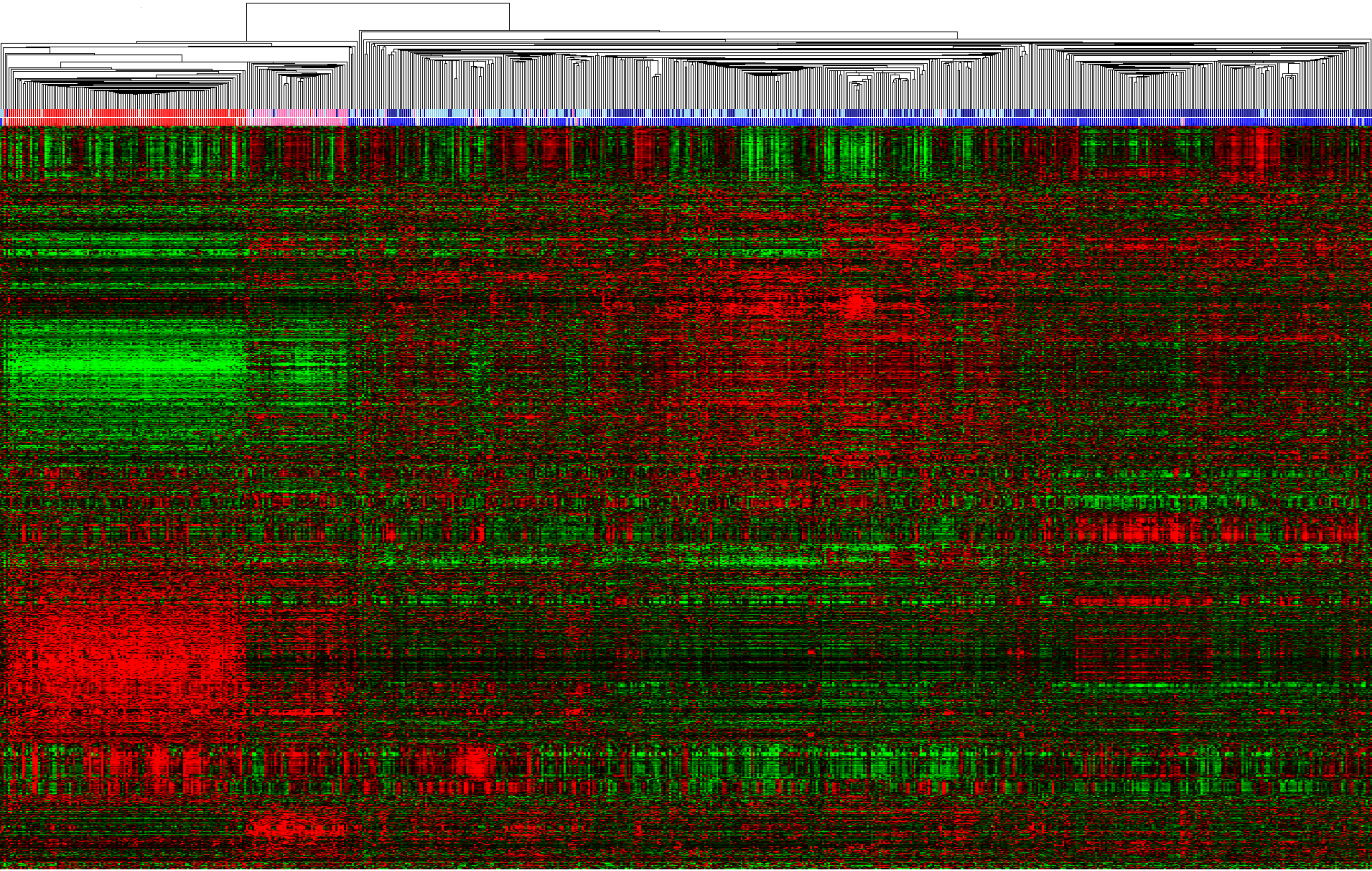
TCGA heatmap. Hierarchical clustering of the 1028 most variable genes in the TCGA dataset. The upper row in the colour bar at the top of the heatmap shows the PAM50 classification (basal, red; HER2, pink; luminal A, dark blue; luminal B, light blue; normal, grey). The lower row in the colour bar at the top of the heatmap shows the LAB classification (basal, red; molecular apocrine, pink; luminal, blue; unknown, grey). To view the individual gene and tumour annotations please use the cdt file in Sup Data 7 or use the script in Sup Data 1 to regenerate the full input files for Cluster.

Sup data 1. Script to classify the tumours and create the figures.

Sup data 2. README file with instructions to run the script.

**Sup data 3. Names of samples and batches in the EORTC data**. Chip type 1 (”PF” names) is Affymetrix U133A; chip type 2 (”HB names”) is Affymetrix X3P. The GSM names are from the NCBI GEO database. The EORTC names are identifiers from the EORTC 10994 clinical trial.

Sup data 4. Annotation file for the U133A chip.

Sup data 5. Annotation file for the X3P chip.

Sup data 6. Treeview cdt file for Sup Fig 1. To regenerate the heatmap, perform correlation (uncentered) centroid hierarchical clustering in Cluster without adjusting the matrix then open the atr, gtr and cdt files in Treeview.

Sup data 7. Treeview cdt file for Sup Fig 2. To regenerate the heatmap, perform correlation (uncentered) centroid hierarchical clustering in Cluster without adjusting the matrix then open the atr, gtr and cdt files in Treeview.

